# Effect of fosfomycin, *Cynara scolymus* extract, deoxynivalenol and their combinations on intestinal health of weaned piglets

**DOI:** 10.1101/323204

**Authors:** Guadalupe Martínez, Susana N. Diéguez, María B. Fernández Paggi, María B. Riccio, Denisa S. Pérez Gaudio, Julieta M. Decundo, Agustina Romanelli, Fabián A. Amanto, María O. Tapia, Alejandro L. Soraci

**Affiliations:** Área Toxicología, Departamento de Fisiopatología, Facultad de Ciencias Veterinarias, Universidad Nacional del Centro de la Provincia de Buenos Aires, Tandil, Buenos Aires, Argentina.; Centro de Investigación Veterinaria de Tandil (CIVETAN), UNCPBA-CICPBA-CONICET, Facultad de Ciencias Veterinarias, Campus Universitario, Tandil, Buenos Aires, Argentina.; Consejo Nacional de Investigaciones Científicas-CONICET, Buenos Aires, Argentina.; Comisión de Investigaciones Científicas de la Provincia de Buenos Aires (CIC-PBA),, Buenos Aires, Argentina.; Área Producción Porcina, Departamento de Producción Animal, Facultad de Ciencias Veterinarias, Universidad Nacional del Centro de la Provincia de Buenos Aires, Tandil, Buenos Aires, Argentina.

## Abstract

Intestinal health of weaning piglets was studied after oral treatments with fosfomycin (FOS), *Cynara scolymus* extract (CSE), deoxynivalenol (DON) and their combinations. Piglets were divided in groups and received different treatments during 15 days, namely DON (1mg/kg of feed), FOS administered into the drinking water (30 mg/kg b.w.), CSE (300 g/ton of feed) and all possible combinations including a control group that received clean balanced diet. At day 15, three piglets from each group were euthanized and gastrointestinal tract samples were immediately taken to evaluate pH, bacteriology (enterobacteria and lactic acid bacteria), volatile fatty acids concentration (VFAs), disaccharidases activity (lactase, sucrase and maltase), histology (intestinal absorptive area [IAA] and goblet cells count) and adherence of bacteria to intestinal mucus. Animals receiving FOS and CSE treatments exhibited evident beneficial intestinal effects compared to animals receiving diets free from these compounds. This was revealed by a lower enterobacteria population together with a lower E/L, an enhanced production of butyric acid, an increased enzymatic activity (particularly maltase), and a greater IAA and goblet cells count along with an increase in pathogenic bacteria adherence to intestinal mucus. Interactions between both treatments resulted in similar beneficial effects as their individual administration. On the contrary, DON produced detrimental effects on intestinal health as a decrease was observed on volatile fatty acids production, enzymatic activity and goblet cells count in animals receiving diets containing sub- toxic concentrations of this mycotoxin. The knowledge of the intestinal effects of these compounds contributes to understand the physiological and pathological gut changes and their potential productive consequences.

## Introduction

Weaning is considered one of the most critical periods of pig production because of its highly negative impact on health and productive performance of piglets, mainly in the first post-weaning days. During this period, the animals are exposed to physiological, immunological, microbiological, social, environmental and nutritional factors that lead to post-weaning stress (1–3). In order to overcome this situation a common, though not rational practice, has been the prophylactic use of antibiotics in intensive pig production. Fosfomycin ((cis 1-2 epoxy propyl) phosphonic acid, FOS) is a broad spectrum bactericide antibiotic, widely used in pig farms in Central and South America, South Africa and Southeast Asia. At weaning FOS is indicated for the treatment of several bacterial infections (*Haemophilus parasuis*, *Streptococcus suis*, *Pasteurella multocida*, *Bordetella bronchiseptica*, *Staphylococcus hyicus*, *Escherichia coli*, etc.) associated to stress (4).

In addition, vegetable extracts, particularly *Cynara scolymus* extract (CSE), have long been used in different species for their hepatoprotective and digestive roles, exerting a choleretic- cholagogue effect, increasing bile concentrations at small intestine level and thus enhancing fat and lipophilic vitamins absorption. In animal production, these compounds are used as feed additives to improve zootechnical parameters (5–8) and they have shown further beneficial consequences on intestine and liver functions. Nowadays enteroprotective, trophic, antitoxic and antimicrobial effects are ascribable to bile action (9–12). CSE is used in intensive pig and avian productions. It is obtained from the leaves of the plant and contains caffeolquinic acid derivatives which are known for their choleretic- cholagogue effect in different species (7,13,14), including pigs (15).

Among weaning stress factors, the presence of anti-nutritional compounds in feed, such as mycotoxins, negatively influences the productive performance of animals. Deoxynivalenol (DON) is a mycotoxin produced by *Fusarium* species, being pigs the most susceptible species to its toxic effects (16,17). Formerly DON was also called vomitoxin, referring to its emetic effect (18,19). Other clinical signs that have been described include reduction in feed intake and complete feed refusal, immunosuppression, haemorrhage and eventually, circulatory shock (20–22). However there is little information on the possible subclinical effects associated to the ingestion of feed contaminated with low DON concentrations, which is highly likely to occur (19,23,24).

In the productive reality, in innumerable situations, but mostly during weaning, antibiotics, natural extracts and mycotoxins coexist in the animals´ diet, and consequently in the gut, regardless the potential interactions among them.

The aim of this study was to evaluate the effect of FOS, CSE, DON and their interactions on the intestinal health of weaning piglets.

## Materials and methods

### Animals

The study was carried out according to guidelines of the Animal Welfare Committee of the Faculty of Veterinary Sciences UNCPBA, Argentina, for animal handling and experimentation. One hundred and sixty, healthy, 21 days old weaned piglets (6.26 ± 0.4 kg body weight [b.w.]) of the same genetic line from a commercial farm were used. Piglets were housed in an environmentally controlled barn (22±5°C; light: dark cycle 12:12 h; relative humidity 45-65%), given free access to feed (commercial feed: 3.0 Kcal/Kg of metabolizable energy) and water, and were checked daily.

### Antibiotic, natural extract and mycotoxin

*Fosfomycin (FOS):* Calcium fosfomycin was provided by Bedson S.A. laboratory (Fosbac^®^, Pilar, Buenos Aires, Argentina). The antibiotic dose was 30 mg/kg b.w. administered via drinking water. Water consumption was measured by a water flow meter installed at the entrance pipeline of the weaning room two days before the beginning of the trial. Medicated water was prepared daily at 8.00 am, considering water consumption and mean piglets weight.

*Cynara scolymus extract (CSE):* This natural extract was provided by Bedson
S.A. laboratory (Bedgen40^®^, Pilar, Buenos Aires, Argentina). Three hundred grams of CSE were uniformly mixed with one ton of feed (15 mg/kg b.w.).

*Deoxynivalenol (DON):* The mycotoxin was produced in our laboratory by growing *Fusarium graminearum* NRRL 28063 in corn at 25°C for 25 days. For DON quantification, samples of ground corn were extracted twice with water/acetonitrile and then with hexane by liquid-liquid extraction. Extracts were passed through DONPREP columns (R-Biopharm, Acre Road, Glascow, Scotland) and evaporated to dryness at 40°C. The dry extract was reconstituted with MilliQ water and filtered through 0.22 μm nylon membranes before Injection into HPLC UV/VIS for quantification. A Gilson HPLC system equipped with a Gilson 151 UV-Vis detector and Gilson 712 software was used for data analysis (Gilson, Inc., Middleton, USA). The column was a C18; 250 mm × 3.00 mm Sinergy Hydro RP 4 μm (Phenomenex, Torrance, United States) maintained in at 35°C. The mobile phase was water: acetonitrile (90:10) at 0.5 ml/min flow rate. DON was detected at 222 nm and its retention time was 8.7 min.

Convenient aliquots of ground contaminated corn were uniformly mixed with feed in order to obtain 1 mg DON/ Kg (50 μg/kg b.w.).

### Experimental groups

Weaning piglets were randomly assigned to one of eight groups, which were subjected to different treatments for a 15 days period. The dietary treatments were as follows: A) balanced diet containing DON (1mg/kg of feed), B) balanced diet and FOS administered into the drinking water (30 mg/kg b.w.), C) balanced diet containing CSE (300 g/ton of feed), D) balanced diet containing DON (1mg/kg of feed) and FOS (30 mg/kg b.w.) into the drinking water, E) balanced diet containing DON (1mg/kg of feed) plus CSE (300 g/ton of feed), F) balanced diet containing CSE (300 g/ton of feed) and FOS (30 mg/kg b.w.) into the drinking water, G) balanced diet containing DON (1mg/kg of feed) plus CSE (300 g/ton of feed) and FOS (30 mg/kg b.w.) into the drinking water and, H) balanced diet without FOS, CSE or DON.

After 15 days of treatment, three piglets of each group were randomly selected and euthanized for sampling of the gastrointestinal tract.

### pH determination

As soon as each sample was obtained, pH was measured with a pH meter (UP-25, Denver Instrument Company, Denver, Colorado, EE. UU.) in the following portions of the gastrointestinal tract: caudal portion of the stomach, ileum (15 cm proximal to ileocaecal valve), caecum and colon (20 cm distal from caecum).

### Enterobacteriaceae/Lactic acid bacteria ratio (E/L)

The E/L has traditionally been used to determine balance of intestinal microbiota in pigs (25). It has been demonstrated that a greater resistance to gastrointestinal diseases is acquired when animals show a lower E/L (26–28).

The intestinal contents from ileum (15 cm proximal to the ileocaecal valve), caecum and colon (20 cm distal from caecum) were collected and kept at 4°C until arrival to the laboratory. One g of sample was diluted in 9 ml of peptone water and homogenized by continuous agitation. Counting of viable bacteria was performed by plating serial 10-fold dilutions (in 1% peptone water) onto MRS agar (Britania S.A.) for *Lactic acid bacteria* (LAB) representative of beneficial bacteria in pigs, and onto Mac Conkey agar (Britania S.A.) for *Enterobacteriaceae* representative of commensal Gram negative bacteria (29–32). Colonies were counted, log transformed and expressed as colony forming units per gram of digesta (CFU/g).

### Volatile fatty acids (VFAs)

The caecal content was immediately diluted with phosphoric acid (in a 4:1 proportion) for preservation and kept at −70°C until analyzed. Concentrations of VFAs were determined using gas liquid chromatography according to the method described by Jouany (33). A Shimadzu chromatograph (Model GC-17A, Kyoto, Japan) with a 19091N-133 Innowax 30M column (Agilent, Santa Clara, CA, USA) was used. A mixture of 10 mM Supelco VFAs (C2 to C10) and 2-ethyl-butyric acid (Fluka) as internal standard were used to build calibration curves.

### Disaccharidases activity

The digestive function of the intestine can be evaluated by the activity of disaccharidases present in the microvilli or brush border of the enterocytes (34,35). The evaluation of these enzymes gives information on the physio pathological status of the intestinal mucosa (36).

The four portions of the small intestine (duodenum, proximal jejunum, medium jejunum and ileum) were opened along the mesenteric border and washed with saline solution to eliminate the mucus and remaining intestinal contents. The mucosa was scraped off with a scalpel and 1.000 g of this material was weighed. Then, saline solution (2 ml) was added to the intestinal mucosa and it was ground with a dispersing instrument (Ultra-Turrax^®^) and a Potter homogenizer. Samples were then cold-centrifuged at 4°C and 6630 rpm for 10 min. The supernatant was used as crude enzyme solution and it was stored at − 20°C until analysis. The protein concentration of each homogenate was determined by Bradford method using bovine serum albumin as standard (37). The activity of sucrase, lactase and maltase was determined by quantification of released glucose, according to Dahlqvist method (38). Briefly, the homogenate supernatants were diluted, added to an equal volume of 0.1 M sodium maleate buffer (pH 6.0) containing 56 mM lactose, sucrose or maltose, and incubated for 1 h at 37°C. Then, the mixtures were added to the glucose oxidase-peroxidase reagents (Sigma Chemical Company, USA) containing O-dianisidine as chromogen. The absorbance was measured using a spectrophotometer (Dupont, Sorvall Instruments) at 450 nm. The activity of disaccharidases was expressed as U/mg protein. One U is defined as the amount of enzyme that hydrolyses 1 mmol of lactose, sucrose or maltose in 1 min under the standard assay conditions.

### Histological study

Different measures on villi and crypts can be correlated to nutrient absorption capacity, possible structural alterations of intestinal mucosa and consequent productive yield (34,39–47).

Samples of medium jejunum (1.5 m from stomach) and ileum (20 cm proximal to ileocaecal valve) were washed with saline solution to remove the intestinal content, transversally cut and fixed in Bouin solution (75% saturated picric acid, 20% formaldehyde and 5% acetic acid). After 24 h of fixation, the samples were embedded in paraffin and stained with haemotoxylin and eosin (H&E) and periodic acid-Schiff (PAS).

The intestinal mucosa was examined under light microscope and measured by the Image Analysis Software (ToupTek™ ToupView™). The length of villi and width of villi and crypts were measured in H&E- stained sections. The goblet cells count in villi and crypts (expressed as goblet cells/ 100 villi or crypts) was determined using PAS staining (48,49). Means were calculated for each group. The mathematical model proposed by Kisielinski *et al* was used to estimate the intestinal absorptive area (IAA) using the following equation (50):

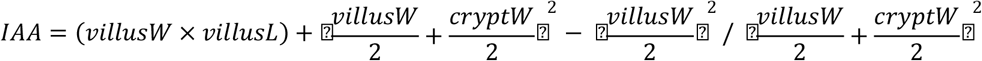

being, *IAA*= intestinal absorptive surface area, *villusW*= villi mean width, *villusL*= villi mean length, and *cryptW*= crypts mean width.

Goblet cells count was used as index of the secretory capacity and the production of protective intestinal mucus (51).

### Adherence of bacteria to intestinal mucus

Mucus quality has been evaluated by its ability to adhere *E. coli*, since bacterial adhesion is associated with the protective and antimicrobial functions of mucus favoring bacterial elimination by the rapid removal of mucus by peristaltic movements (52,53). The interaction between the glycoproteins of the outer layer of the intestinal mucus and *E. coli* would prevent the attachment of bacteria to epithelial cells and subsequent damage (51,53–58).

Ileum samples (15 cm proximal to ileocaecal valve) were opened along the mesenteric border. The mucus was carefully scraped off with a scalpel (leaving intestinal mucosa intact), collected into sterile tubes and kept at −70°C until analyzed.

The adherence of bacteria to the intestinal mucus was analyzed according to Bai *et a.* (59). One hundred milligrams of mucus were diluted with 1.5 ml of saline solution and centrifuged (12.000 rpm, 10 min, 4°C) to remove cell debris and bacteria. The supernatant was sterilized by filtration (13 mm × 0.22 μm nylon filter membranes) and the filtered solution was defined as the original crude mucus that contained glycoproteins responsible for bacteria adherence. A concentration of 10^3^ CFU/ml of *Escherichia coli* O157:H7 was incubated with supernatant containing crude mucus for 30 min, at 37°C under continuous agitation. Then the tubes were centrifuged (12.000 rpm, 10 min, 4°C) and pellets (with adhered and not adhered bacteria) were resuspended in 400 μl saline solution and further centrifuged (2000 rpm, 4°C, 2 min). Two fractions were obtained, the pellet which contained adhered bacteria and the supernatant which contained not adhered bacteria. Aliquots from pellet and supernatant were spread on Mac Conkey Agar with Sorbitol (Britania S.A.) and incubated under aerobic condition for 24 h at 37 °C for colonies count. Results were expressed as percentage of adhered bacteria to the intestinal mucus.

### Statistical analyses

A 2×2×2 factorial arrangement was used to evaluate interactions between FOS (0 *vs*. 30 mg/kg b.w.), CSE (0 *vs*. 300 g/ton of feed) and DON (0 *vs*. 1 mg/kg feed) on the intestinal health of weaned piglets. The response variables (pH, intestinal bacteria, VFAs, dissacharidases activity, IAA, goblet cells and percentage of adhered bacteria to intestinal mucus) were subjected to analysis of variance (ANOVA) by GLM procedure of SAS V9.3 (SAS Institute Inc., Cary, NC, USA). Differences between treatments were declared significant when *p* < 0.05. When significant interactions were observed, contrasts were used to compare the different levels of each treatment. Data are presented in tables as means and mean standard error (SEM).

## Results

### pH

pH values are shown in Table 1. No statically significant differences on pH were found neither in gastrointestinal (GI) portions studied for groups treated with CSE and DON nor in interactions between the different factors. In FOS treated groups, no statically significant effects were found in caudal portion of stomach and ileum, but piglets that received FOS showed a lower pH (*p*< 0.01) in caecum and colon. The mean caecal pH was 5.51±0.33 in FOS treated groups and 6.90±0.29 in FOS free groups. The mean pH in the colon was 6.21±0.30 in FOS treated groups and 7.36±0.26 in FOS free groups.

**Table 1.**
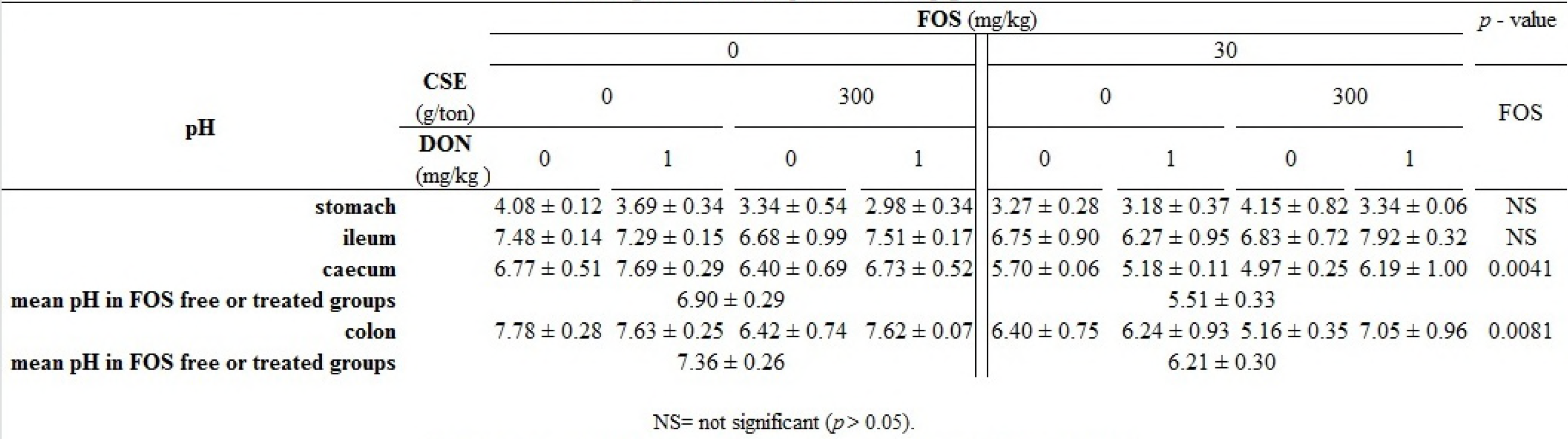
Effect of fosfomycin (FOS), *C. scolymus* extract (CSE), deoxynivalenol (DON) and their combinations on the gastrointestinal pH of weaned piglets1

### Enterobacteriaceae, lactic acid bacteria and E/L

There was no effect of none of the treatments on the studied bacteria at ileum level. LAB counts from caecum and colon did not show any significant differences among treatments and effects of DON on *Enterobacteriaceae* in these intestinal portions were neither detected. In caecum and colon, FOS and CSE treated groups showed lower *Enterobacteriaceae* population and E/L regardless the presence of DON (Table 2). A significant antagonistic interaction was observed between FOS and CSE on *Enterobacteriaceae* count (*p*= 0.0004) and consequently on the E/L (*p*= 0.0016) at caecum level. In this case, the effect of both treatments was less pronounced than the effect they produced as individual factors. An indifferent interaction was observed for *Enterobacteriaceae* count (*p*= 0.0004) and E/L (*p*= 0.0114) at colon level when FOS and CSE were combined, i.e., the effect produced by the combination of FOS and CSE was similar to the one observed when they were administered individually.

**Table 2.**
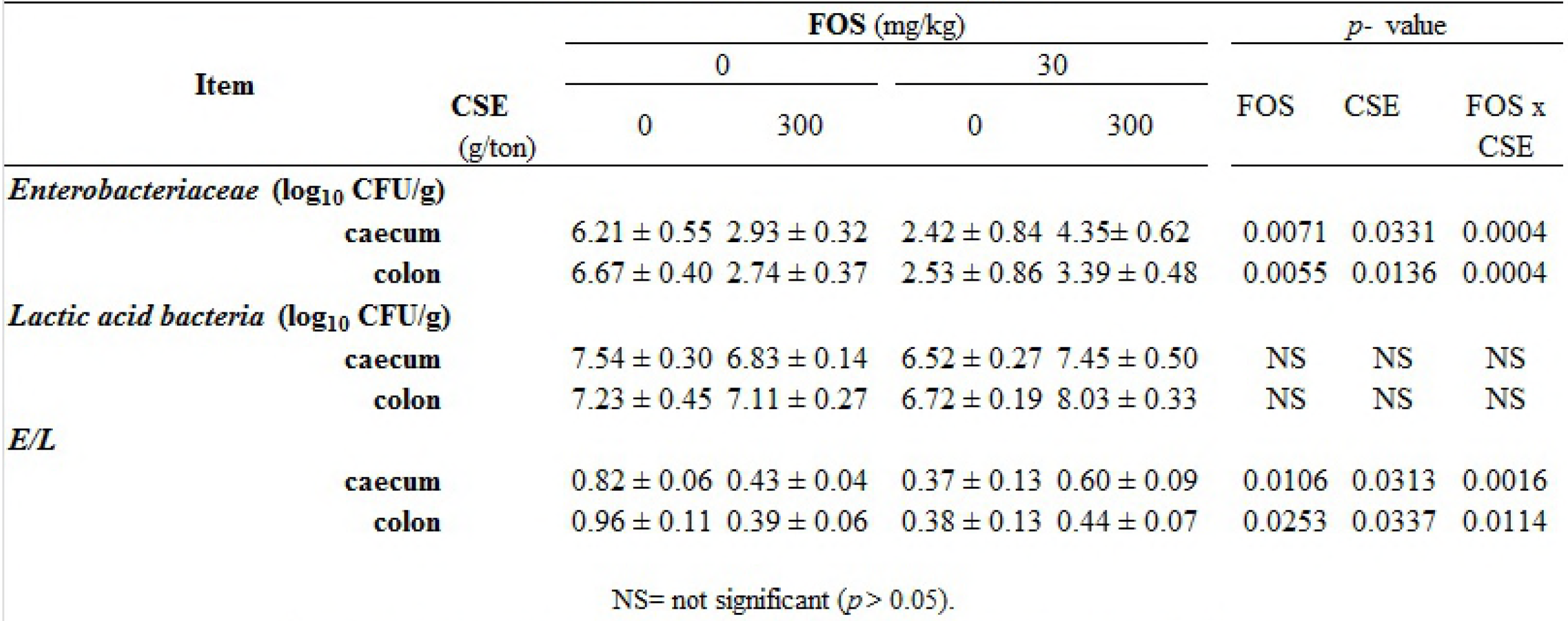
Effect of the interaction between fosmomycin (FOS) and *C. scolymus* extract (CSE) on the intestinal bacteria of weaned piglets1

### Volatile fatty acids

Concentrations of VFAs were not modified in FOS treated groups and interactions between different factors were not significant (*p*> 0.05). CSE treated groups increased the concentrations of butyric acid (*p*= 0.033). For DON treated groups lower acetic (*p*= 0.0104) and butyric (*p*= 0.0001) acids and lower total VFAs concentrations (*p*= 0.0021) were detected (Table 3).

**Table 3.** Effect of fosfomycin (FOS), *C. scolymus* extract (CSE), deoxynivalenol (DON) on VFAs (momol/L) in the caecum of weaned piglets1

| VFAs | FOS (mg/kg) |  |  | CSE (g/ton) |  |  | DON (mg/kg) |  |  |
| --- | --- | --- | --- | --- | --- | --- | --- | --- | --- |
|  | 0 | 30 | p - value | 0 | 300 | p - value | 0 | 1 | p - value |
| Acetic acid | 49.71 ± 4.5 | 60.35 ± 7.31 | NS | 50.29 ± 4.75 | 58.77 ± 6.88 | NS | 62.35 ± 5.55 | 43.01 ± 4.33 | 0.0104 |
| Propionic acid | 13.62 ± 1.27 | 16.44 ± 1.60 | NS | 13.95 ± 1.30 | 15.79 ± 1.60 | NS | 17.74 ± 1.10 | 10.84 ± 1.16 | NS |
| Butyric acid | 5.87 ± 0.60 | 6.78 ± 1.03 | NS | 5.83 ± 0.65 | 6.76 ± 0.93 | 0.033 | 8.00 ± 0.65 | 3.92 ± 0.36 | 0.0001 |
| Total VFAs | 71.49 ± 6.42 | 85.53 ± 9.69 | NS | 72.55 ± 6.72 | 83.07 ± 9.25 | NS | 90.76 ± 7.14 | 59.26 ± 5.81 | 0.0021 |

NS= not significant ( $p > 0.05$ ).
<sup>1</sup> Interaction between the different factors was not significant ( $p > 0.05$ ).

### Disaccharidases activity

There were not significant interactions between FOS, CSE and DON on disaccharidases activity *(p*> 0.05).

It was found that the activity of maltase in the different intestinal regions in piglets from FOS treated groups was significantly higher (*p*< 0.05) than that observed in FOS free groups. FOS treatments also increased sucrose and lactase activity in proximal and medium jejunum and ileum though this effect was not statistically significant. Treatments with CSE produced higher maltase activity in ileum (p= 0.0020). However, an effect on the activity of sucrase and lactase was not observed. DON showed negative effects for all enzymes in all intestinal portions, being enzymatic activity lower for pigs fed diets supplemented with DON when compared to those without DON supplementation. P value< 0.05 was observed for maltase and lactase activity in duodenum and proximal jejunum, sucrase and lactase in medium jejunum and maltase in the ileum (Table 4).

**Table 4.** Effect of fosmomycin (FOS), *C. scolymus* extract (CSE), deoxynivalenol (DON) on disaccharisases acticity.

| Item | FOS (mg/kg) |  |  | CSE (g/ton) |  |  | DON (mg/kg) |  |  |
| --- | --- | --- | --- | --- | --- | --- | --- | --- | --- |
|  | 0 | 30 | p - value | 0 | 300 | p - value | 0 | 30 | p - value |
| **<u>Duodenum</u>** |  |  |  |  |  |  |  |  |  |
| Maltase | 1427.85 ± 169.23 | 2422.91 ± 338.39 | 0.0008 | 1951.87 ± 350.73 | 1862.65 ± 246.43 | NS | 2462.37 ± 339.66 | 1391.43 ± 147.09 | 0.0004 |
| Sucrase | 37.02 ± 15.26 | 36.86 ± 8.85 | NS | 41.32 ± 15.17 | 32.90 ± 10.07 | NS | 41.69 ± 10.17 | 32.56 ± 14.43 | NS |
| Lactase | 86.84 ± 28.70 | 88.19 ± 27.68 | NS | 72.60 ± 18.06 | 101.23 ± 34.11 | NS | 141.09 ± 33.98 | 38.01 ± 8.36 | 0.0092 |
| **<u>Proximal jejunum</u>** |  |  |  |  |  |  |  |  |  |
| Maltase | 1964.33 ± 232.52 | 3419.02 ± 456.61 | 0.0028 | 2727.34 ± 465.01 | 2602.81 ± 362.69 | NS | 3420.65 ± 461.72 | 1962.83 ± 223.01 | 0.0028 |
| Sucrase | 89.26 ± 29.52 | 133.11 ± 51.55 | NS | 133.87 ± 28.64 | 88.55 ± 49.20 | NS | 145.62 ± 53.67 | 77.71 ± 23.98 | NS |
| Lactase | 162.66 ± 49.17 | 257.84 ± 91.37 | NS | 214.62 ± 55.72 | 202.56 ± 84.92 | NS | 324.52 ± 93.05 | 101.12 ± 22.84 | 0.041 |
| **<u>Medium jejunum</u>** |  |  |  |  |  |  |  |  |  |
| Maltase | 2712.30 ± 293.58 | 4333.48 ± 493.03 | 0.0064 | 3854.78 ± 549.21 | 3154.17 ± 349.49 | NS | 4038.04 ± 432.94 | 2985.01 ± 440.51 | NS |
| Sucrase | 164.87 ± 48.53 | 284.39 ± 66.33 | NS | 259.65 ± 63.40 | 187.70 ± 55.14 | NS | 319.49 ± 69.41 | 132.47 ± 34.25 | 0.0147 |
| Lactase | 201.64 ± 62.08 | 342.47 ± 79.00 | NS | 316.65 ± 72.98 | 225.47 ± 71.34 | NS | 407.83 ± 80.99 | 141.30 ± 39.20 | 0.0105 |
| **<u>Ileum</u>** |  |  |  |  |  |  |  |  |  |
| Maltase | 1047.08 ± 129.29 | 1464.89 ± 265.51 | 0.0337 | 947.73 ± 122.61 | 1524.46 ± 239.42 | 0.002 | 1623.23 ± 259.01 | 900.92 ± 75.58 | 0.0007 |
| Sucrase | 14.20 ± 9.13 | 59.01 ± 45.49 | NS | 15.72 ± 9.81 | 54.16 ± 42.12 | NS | 57.68 ± 45.56 | 15.42 ± 9.34 | NS |
| Lactase | 10.52 ± 3.42 | 18.26 ± 5.51 | NS | 14.16 ± 4.36 | 14.30 ± 4.87 | NS | 15.69 ± 4.86 | 12.89 ± 4.42 | NS |

NS= not significant ( $p > 0.05$ ).<sup>1</sup> Interaction between the different factors was not significant ( $p > 0.05$ ).

### Intestinal absorptive area and goblet cells

There was an evident increase in the IAA of medium jejunum in the presence of FOS, CSE and the combination of both factors (*p*< 0.05). The co-administration of FOS and CSE showed an indifferent type interaction at this level. IAA of ileum increased in piglets that received CSE and an antagonistic interaction between FOS and CSE was detected (*p*< 0.05). The IAA of medium jejunum and ileum was not affected by the treatments with DON (*p*> 0.05), (Table 5).

**Table 5.**
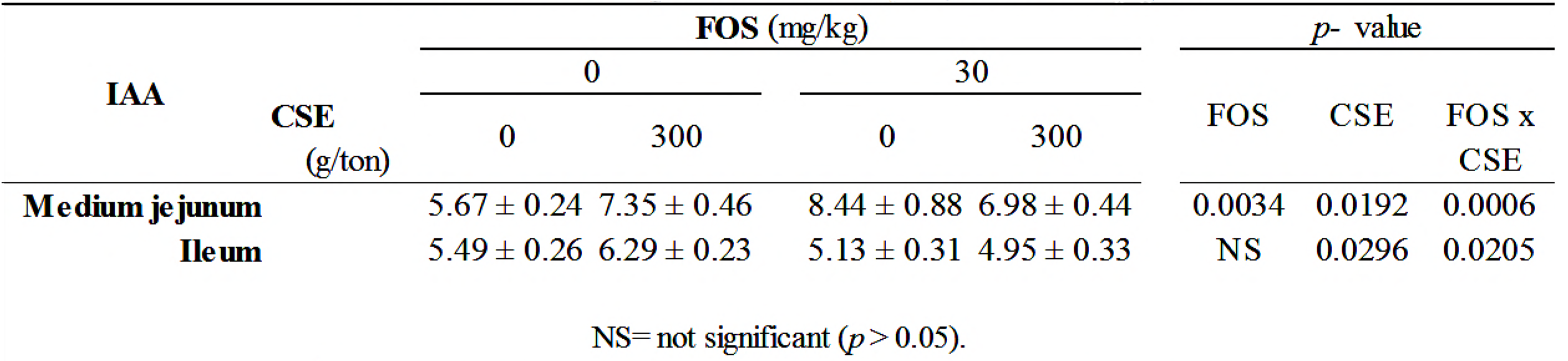
Effect of the interaction between fosfmomycin (FOS) and *C. scolymus* extract (CSE), on the intestial absorptive area (IAA, μm^2^) in weaned piglets1

Generally, the number of intestinal goblet cells increased with FOS and CSE treatments, whereas a decrease was evident in goblet cells from villi after DON treatments. Goblet cells count in crypts of ileum increased in FOS treated groups (*p*= 0120). The treatments with CSE increased the count of these cells in villi (*p*= 0.0159) and crypts (*p*= 0.0143) of medium jejunum. A negative effect of DON was observed in goblet cells count in villi of medium jejunum (*p*= 0.0125) and ileum (*p*= 0.0336) (Table 6). No significant interactions were detected between FOS, CSE and DON on goblet cells count (*p*> 0.05).

**Table 6.** Effect of fosfomycin (FOS), *C. scolymus* extract (CSE), deoxynivalenol (DON) on globlet cells/100 villi and goblet cells/100 crypts in the small intestine of weaned piglets1

| Item | FOS (mg/kg) |  | *p* - value | CSE (g/ton) |  | *p* - value | DON (mg/kg) |  | *p* - value |
| --- | --- | --- | --- | --- | --- | --- | --- | --- | --- |
|  | 0 | 30 |  | 0 | 300 |  | 0 | 30 |  |
| **<u>Medium jejunum</u>** |  |  |  |  |  |  |  |  |  |
| villi | 871.69 ± 84.47 | 1153.33 ± 120.80 | NS | 886.47 ± 107.87 | 1114.62 ± 93.93 | 0.0159 | 1087.80 ± 103.30 | 882.31 ± 102.77 | 0.0125 |
| crypts | 1155.69 ± 78.55 | 1291.67 ± 86.14 | NS | 1095.73 ± 70.53 | 1350.38 ± 83.63 | 0.0143 | 1221.07 ± 82.77 | 1205.77 ± 85.74 | NS |
| **<u>Ileum</u>** |  |  |  |  |  |  |  |  |  |
| villi | 1045.94 ± 72.8 | 954.58 ± 75.52 | NS | 947.33 ± 81.14 | 1075.38 ± 61.25 | NS | 1102.67 ± 75.37 | 896.15 ± 61.61 | 0.0336 |
| crypts | 1421.31 ± 94.7 | 1677.08 ± 81.43 | 0.012 | 1437.73 ± 103.07 | 1638.46 ± 78.63 | NS | 1526.73 ± 75.23 | 1535.77 ± 121.08 | NS |

NS= not significant ( $p > 0.05$ ).
<sup>1</sup> Interaction between different factors was not significant ( $p > 0.05$ ).

### Adherence of bacteria to the intestinal mucus

Treatments with FOS, CSE and the combination of both resulted in a statistically significant increase in the percentages of adhesion of bacteria to intestinal mucus (*p*< 0.001, *p*= 0.0133 and *p*= 0.0049, respectively) compared to FOS and CSE free groups. In the latter, the adhesion percentage of *E. coli* was 45.71%, whereas FOS or CSE treated groups increased the percentage of adhesion to 83.67% and 72.75%, respectively. The combination of treatments evidenced an indifferent type interaction. In this case, the adhesion percentage of bacteria was 81.61%. The percentage of bacteria adhered to intestinal mucus was not affected by treatments with DON (*p*> 0.05), (Table 7).

**Table 7.**
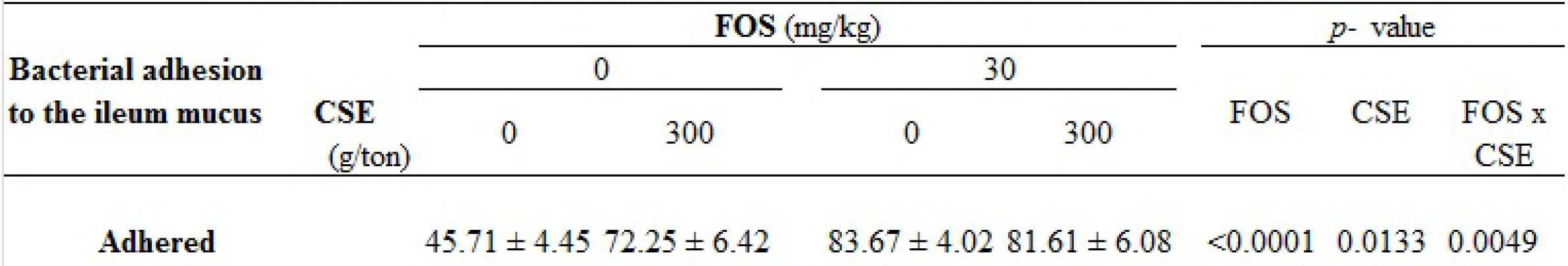
Effect of the interaction between fosfomycin (FOS), *C. scolymus* extract (CSE), on the percentages (%) of adherence of bacteria to intestinal muscus of weaned piglets1

## Discussion

FOS, CSE and DON are commonly found together in the weaning diet. These compounds, individually or combined, may impact on the important morphological, histological and microbiota modifications produced during weaning, affecting the animals´ productive outcome.

### Bacteria, VFAs and pH

LAB populations were not affected by none of the treatments in any of the intestine portions studied. Natural resistance of LAB strains to antibiotics and bile salts, increased by CSE consumption, has been largely demonstrated (17,60–62), The influence of mycotoxins on intestinal microbiota of pigs have been poorly investigated. Available data on the interaction of mycotoxins with bacteria are mainly related to the ability of the intestinal microbiota to detoxify mycotoxins (63–67,24). Results obtained in a study conducted by Waché *et al.* showed that cultivable bacteria diversity in fecal samples was conserved in animals that consumed feed naturally contaminated with DON (2.8 mg/kg) (24). Accordingly, in our study, when piglets received diets containing DON at 1mg/kg, alone or in combination with the other factors, significant changes in CFU counts were neither observed for LAB nor for *Enterobacteriaciae*. In addition, pH values were conserved in all gastrointestinal tract portions after DON treatments. It is likely that gut bacteria possess resistance mechanisms against this mycotoxin, in fact *in vitro* studies identified intestinal bacterial strains that promote metabolism, binding or detoxification of DON (64,67). By contrast, VFAs concentrations were lowered. The normal concentration of VFAs in the caecum varies according to the content and composition of the raw material in the diet, being around 80 mmol/L for this stage of pig rearing (68–70). In the present study, the decrease in VFAs at caecum level, where the mycotoxin is metabolized, could be explained by a detrimental effect of DON on the metabolism of culture independent bacterial populations as it has been previously demonstrated (17,24).

A lower count of *Enterobacteriaciae* population and E/L in caecum and colon was observed in pigs treated with CSE. It has been recently demonstrated by our research group that using CSE as feed additive substantially increases bile production in pigs (15). Important bile effects on the intestinal microbiota have been described involving two main mechanisms: direct detergent action on bacterial cell membranes (mainly in proximal intestine) and an indirect action by interacting with specific nuclear receptors (FXR, TGR 5, mainly in large intestine) and thus inducing antimicrobial peptides synthesis (10,71–73). Furthermore, Cremers *et al.* indicated that bile acid salts have profound effects on many key proteins in bacteria (74). Results from different studies suggest that bile salts could potentially induce DNA damage through oxidative stress in *E. coli* (75–79). Therefore bile acids are thought to have destructive effects on gut microbes except for some bile acid tolerant bacteria. LAB can tolerate biliary acids by expressing bile salts hydrolases (80). This might have contributed to lower *Enterobacteriaceae* count in CSE treated groups in our study without altering LAB.

The administration of CSE in the piglets´ diet increased the concentration of butyric acid, an important energetic VFA in large intestine (81–84). This finding is in agreement with other scientific studies that detected an increase in the proportion of butyrate and an equal or lower concentration of acetate in diets containing other natural extracts (85–90). The no significant change in the levels of acetate could be attributed to the fact that butyrate producing bacteria are able to use acetate as a substrate. In this way, acetic acid constitutes a type of substrate for cross-feeding interactions that occur among colonic bacteria (91–95).

FOS reduced *Enterobacteriaceae* populations in caecum and colon exerting a bactericide effect, related to its low oral bioavailability (96). LAB populations, capable of resisting relatively high bactericide antibiotic concentrations through different adaptive mechanisms (97,98), were not affected by FOS. Further, a reduction in pH was observed in caecum and colon as consequence of the diminished E/L in FOS treated groups. These findings, together with an increase of butyrate production (*p*> 0.05) represent an important favorable aspect of intestinal health in weaning piglets.

Interactions observed between FOS and CSE treatments could be explained by a possible interference of their mechanisms of action: The modes by which antimicrobial intestinal peptides kill bacteria are varied. The cytoplasmic membrane is a frequent target, but peptides may also interfere with DNA and protein synthesis, protein folding, and cell wall synthesis. Thereby, some peptides form a complex with different cell wall precursors inhibiting cell wall biosynthesis. On the other hand, FOS is transported into bacteria via both glycerol-3-phosphate and hexose phosphate membrane transporter systems. Besides, it interferes with the cytoplasmic step of bacterial cell wall biosynthesis, the formation of the peptidoglycan precursor UDP *N*-acetylmuramic acid (99–103). Interference between the action of FOS and intestinal peptides (induced by biliary acids through nuclear receptors), at cytoplasmatic or membrane transporter level could occur. Moreover, some cytoplasmic peptides show bacteriostatic effects that could antagonize bactericidal effect of FOS that requires bacteria to grow at log phase to exert it´s action; i.e. antagonistic or indifference effects may be due to inhibition of bacterial growth by static agents (104).

### Intestinal morpho-physiology

Clinical symptoms characteristic of DON intoxication were not observed in the animals under study. In order to evaluate intestinal health, morphological and physiological integrity of intestinal mucosa was studied. The mycotoxin DON administered at 1mg/Kg of feed in our experiment did not affect IAA, which is in agreement with studies that indicate that higher DON concentrations are needed to deteriorate the tissue at this level (105). However, in the present study, treatments with DON adversely affected the number of goblet cells. Similarly, Obremski *et al.* obtained a lower goblet cells count in jejunum of piglets maintained on diets contaminated with DON for 14 days (106). In addition, Bracarense *et al.* and Gerez *et al.* also reported lower goblet cell counts in jejunum after administration of 1.5 to 3 mg/kg DON respectively in the diet of animals during 4 to 5 weeks (107,108). Apart from a lower goblet cells count, a lower expression of mucins (mainly MUC1, MUC2 and MUC3) by these cells would be expected after the ingestion of low DON concentrations (109–111). In our study this effect was reflected by a lower adherence of *E. coli* to mucus (p > 0.05; data not shown). Moreover, disaccharidases activity decreased with DON treatments, particularly maltase and sucrose, in the different portions of the intestine. The undesirable effect of the mycotoxin could be a consequence of its mechanism of action as a potent inhibitor of protein synthesis, including the synthesis of disaccharidases (21).

After FOS treatments, IAA and goblet cells were considerably increased. In a previous study, Pérez Gaudio *et al.* demonstrated a protective effect of the antibiotic FOS on in vitro cell cultures that would favor a trophic effect on intestinal mucosa (112). On the other hand, certain antibiotics modulate physiological inflammation decreasing the catabolic cost of maintaining immune response, thereby favoring mucosal anabolic processes (113–117). A greater goblet cells count improved mucus production which was revealed by a greater pathogenic bacterial adhesion. Enzymatic activity was also increased in FOS treated groups, being maltase the most active disaccharidase, as expected for the age and diet of the animals (118).

CSE as an additive in the diet significantly increased IAA and goblet cells. These findings are consistent with previous works which reported that using different sources of natural extracts increased villi height and villi: crypts ratio in the small intestine of weaned piglets (95,119,120). In pigs, the action of bile acids on the G protein-coupled bile acid receptor (TGR5) found in enteroendocrine cells stimulates secretion of glucagon like peptides (GLP)-1 and 2, which function respectively as the major incretin hormone involved in glucose homeostasis and key trophic hormone in intestinal adaptation and growth in response to food ingestion. In fact, the induction of GLP-2 secretion, by TGR 5, is involved in the trophic action of bile acids in the intestinal lumen (121,122). The observed increase in IAA and goblet cells in our study could be explained by the direct trophic effect of the increased bile production (73,123) when CSE is added to the diet. As stated before, the increased bacterial adherence to mucus would be a direct consequence of the increased number of goblet cells rendering a better mucus quality. Maltase activity, which plays an important role at weaning, was increased in CSE treated groups at ileum level. This could be related to the trophic effect of bile acids, augmented after CSE administration, in this portion of the intestine through interaction with specific nuclear receptors (10,12,73,124).

Beneficial effects observed after co-administration of FOS and CSE on IAA and bacterial adherence to mucus did not exceed the benefits of individual treatments (antagonistic or indifferent interactions). It could be possible that anti-inflammatory mechanisms exerted by FOS and biliary acids, that involve cytokines produced by intestine immune cells, interfere at different levels (117,125–135).

## Conclusions

The gastrointestinal mucosa is the first biological barrier that makes contact to different compounds present in feed, and consequently, it could be exposed to dietary toxins. Thereby the intestinal epithelial cells are target for antibiotic, natural extracts used as additives and mycotoxins.

In the present study, we have demonstrated the impact of FOS, CSE and DON on intestinal health parameters. DON showed a deleterious effect at different levels of the intestinal epithelium at sub- toxic concentrations. This could represent a predisposing factor to progressive weight loss, digestive problems and diarrhea as well as a reduction in the intestinal barrier function.

The antibiotic FOS and CSE improved all studied parameters in relation with the intestinal health. Interactions between both treatments resulted in similar beneficial effects as the individual administration, there remains work to be done investigating the specific mechanisms which contribute to this type of interactions.

Finally, the knowledge of the intestinal effects of these compounds contributes to understand the physiological/physio-pathological gut changes and their potential productive consequences. Particularly, CSE could be considered as a nutritional strategy to prevent enteric disorders and improve intestinal health in post-weaned piglets, emerging as a possible alternative to preventive use of antibiotics. In addition, the presence of mycotoxins in feed even at sub-toxic concentrations may cause detrimental gastrointestinal effects and should not be underestimated.

## Acknowledgements

This work was supported by Consejo Nacional de Investigaciones Científicas y Técnicas (PICT 2012- 2398) from Argentina.

The authors would like to thank Edgardo Rodriguez and Sandra E. Pérez for collaborating with this study.

## Footnotes

1 No signeficant effect of CSE and DON on the gastrointestinal pH were detected. Interaction between the different factors was not significant(*p*> 0.05).

1 Effect of mycotoxin treatments on the intestial backteria were not significant.

1 Interaction between the different factors was not significant(*p*> 0.05).

1 The effect of the mycotoxin treatments on the IAA were not significant(*p*> 0.05).

1 Interaction between different factors was not significant(*p*> 0.05).

1 Mycotoxin treatments produced on the adherence of bacteria to intestial mucus.

## References

1. Hampson DJ, Kidder DE. Influence of creep feeding and weaning on brush border enzyme activities in the piglet small intestine. Res Vet Sci. 1986;40(1): 24–31.

2. Lallès J, Bosi P, Smidt H, Stokes CR. Weaning — A challenge to gut physiologists. Livest Sci. 2007;108: 82–93.

3. Heo JM, Opapeju FO, Pluske JR, Kim JC, Hampson DJ, Nyachoti CM. Gastrointestinal health and function in weaned pigs: a review of feeding strategies to control post-weaning diarrhoea without using in-feed antimicrobial compounds. J Anim Physiol Anim Nutr. 2013;97: 207–237.

4. Martineau G-P, Morvan H. Maladies d’élevage des porcs: diagnostics, causes, traitements. France Agricole Editions; 2010.

5. Schiavone A, Righi F, Quarantelli A, Bruni R, Serventi P, Fusari A. Use of Silybum marianum fruit extract in broiler chicken nutrition: influence on performance and meat quality. 2007;91: 256–262.

6. Abbasi F, Samadi F. Effect of Different Levels of Artichoke (Cynara scolymus L.) Leaf Powder on the Performance and Meat Quality of Japanese Quail. Poult Sci J. 2014;2(2): 95–111.

7. Martínez D, Uculmana C. Artichoke extract (Cynara scolymus L.): experiences of use in animal production markets and opportunities for its production in Peru. Agroindustrial Sci. 2016;1: 155–161.

8. Saeed M, Babazadeh D, Arif M, Arain MA, Bhutto ZA, Shar AH, et al. Silymarin: a potent hepatoprotective agent in poultry industry. Worlds Poult Sci J. 2017;73(3): 483–492.

9. Bertók L. Bile acids in physico-chemical host defence. Pathophysiology. 2004;11(3): 139–145.

10. Inagaki T, Moschetta A, Lee YK, Peng L, Zhao G, Downes M, et al. Regulation of antibacterial defense in the small intestine by the nuclear bile acid receptor. Proc Natl Acad Sci USA. 2006;103(10): 3920–3925.

11. Mikov M, Fawcett J, Kuhajda K, Kevresan S. Pharmacology of Bile Acids and their Derivatives: Absorption Promoters and Therapueutic Agents. Eur J drug. 2006;31(3): 237–251.

12. Chiang JYL. Bile acids: regulation of synthesis. J Lipid Res. 2009;50(10): 1955–1966.

13. Speroni E, Cervellati R, Govoni P, Guizzardi S, Renzulli C, Guerra MC. Efficacy of different Cynara scolymus preparations on liver complaints. J Ethnopharmacol. 2003;86: 203–211.

14. Wegener T, Fintelmann V. Pharmacological properties and therapeutic profile of artichoke (Cynara scolymus L.). Wien Med Wochenschr. 1999;149(8–10): 241–247.

15. Martínez G, Diéguez SN, Rodríguez E, Decundo JM, Romanelli A, Fernández Paggi MB, et al. Effect of Cynara scolymus and Silybum marianum extracts on bile production in pigs. J Appl Anim Res. 2018;46(1): 1059–1063.

16. Rotter BA. Invited review: Toxicology of deoxynivalenol (vomitoxin). J Toxicol Environ Heal Part A. 1996;48(1): 1–34.

17. Piotrowska, K. Śliżewska, A. Nowak, Ł. Zielonka, Z. Żakowska, M. Gajęcka MG. The Effect of Experimental Fusarium Mycotoxicosis on Microbiota Diversity in Porcine Ascending Colon Contents. Toxins (Basel). 2014;6: 2064–2081.

18. Vesonder RF, Ciegler A, Jensen AH. Isolation of the emetic principle from Fusarium-infected corn. Appl Microbiol. 1973;26(6): 1008–1010.

19. Eriksen GS, Pettersson H. Toxicological evaluation of trichothecenes in animal feed. Anim Feed Sci Technol. 2004;114: 205–239.

20. Lawlor PG, Lynch PB. peer reviewed Mycotoxins in pig feeds 2: clinical aspects. Ir Vet J. 2001;54(4): 172–176.

21. Pestka JJ. Deoxynivalenol: Toxicity, mechanisms and animal health risks. Anim Feed Sci Technol. 2007;137: 283–298.

22. Alizadeh A, Braber S, Akbari P, Garssen J, Fink-gremmels J. Deoxynivalenol Impairs Weight Gain and Affects Markers of Gut Health after Low-Dose, ShortTerm Exposure of Growing Pigs. Toxins (Basel). 2015;7: 2071–2095.

23. Roigé MB, Aranguren SM, Riccio MB, Pereyra S, Soraci AL, Tapia MO. Mycobiota and mycotoxins in fermented feed, wheat grains and corn grains in Southeastern Buenos Aires Province, Argentina. Rev Iberoam Micol. 2009;26(4): 233–237.

24. Waché YJ, Valat C, Postollec G, Bougeard S, Burel C, Oswald IP, et al. Impact of deoxynivalenol on the intestinal microflora of pigs. Int J Mol Sci. 2009;10(1): 1–17.

25. Castillo M, Martín-Orúe SM, Anguita M, Pérez JF, Gasa J. Adaptation of gut microbiota to corn physical structure and different types of dietary fibre. Livest Sci. 2007;109(1–3): 149–152.

26. Han KS, Balan P, Molist Gasa F, Boland M. Green kiwifruit modulates the colonic microbiota in growing pigs. Lett Appl Microbiol. 2011;52(4): 379–385.

27. O’Shea CJ, Sweeney T, Lynch MB, Callan JJ, O’Doherty J V. Modification of selected bacteria and markers of protein fermentation in the distal gastrointestinal tract of pigs upon consumption of chitosan is accompanied by heightened manure odor emissions. J Anim Sci. 2011;89(5): 1366–1375.

28. Chen K, Gao J, Li J, Huang Y, Luo X, Zhang T. Effects of probiotics and antibiotics on diversity and structure of intestinal microflora in broiler chickens. African J Microbiol Res. 2012;6(37): 6612–6617.

29. Macconkey AT. Note on a New Medium for the Growth and Differentiation of the Bacillus Coli Communis and the Bacillus Typhi Abdominalis. Lancet. 1900;156(4010): 20.

30. De Man JC, Rogosa M, Sharpe ME. A medium used for the cultivation of Lactobacilli. J Appl Bacteriol. 1960;23: 130–135.

31. White LA, Newman MC, Cromwell GL, Lindemann MD. Brewers dried yeast as a source of mannan oligosaccharides for weanling pigs. J Anim Sci. 2002;80(10): 2619–2628.

32. Mikkelsen LL, Jensen BB. Effect of fructo-oligosaccharides and transgalacto-oligosaccharides on microbial populations and microbial activity in the gastrointestinal tract of piglets post-weaning. Anim Feed Sci Technol. 2004;117(1–2): 107–119.

33. Jouany JP. Volatlile fatty acid and alcohol determination in digestive contents, silage juices, bacterial cultures and anaerobic fermentor contents. Sci Aliments. 1982; 2:131–144.

34. Pluske JR, Thompson MJ, Atwood CS, Bird PH, Williams IH, Hartmann PE. Maintenance of villus height and crypt depth, and enhancement of disaccharide digestion and monosaccharide absorption, in piglets fed on cows’ whole milk after weaning. Br J Nutr. 1996;76: 409–422.

35. Awad WA, Ghareeb K, Paßlack N, Zentek J. Dietary inulin alters the intestinal absorptive and barrier function of piglet intestine after weaning. Res Vet Sci. 2013;95(1): 249–254.

36. Solaymani-Mohammadi S, Singer SM. Host Immunity and Pathogen Strain Contribute to Intestinal Disaccharidase Impairment following Gut Infection. J Immunol. 2011;187(7): 3769–3775.

37. Bradford MM. A rapid and sensitive method for the quantitation of microgram quantities of protein utilizing the principle of protein-dye binding. Anal Biochem. 1976;72(1–2): 248–254.

38. Dahlqvist A. Method for Assay of Intestinal Disaccharidases. Anal Biochem 1964;7: 18–25.

39. Grant AL, Thomas JW, King KJ, Liesman JS. Effects of dietary amines on the small intestinal variables in neonatal pigs fed soy protein isolate. J Anim Sci. 1990;68: 363–371.

40. Pluske JR, Williams IH, Aherne FX. Villous height and crypt depth in piglets in response to increases in the intake of cows’ milk after weaning. Anim Sci. 1996;62(1): 145–158.

41. van Beers-Schreurs HM, Nabuurs MJ, Vellenga L, Kalsbeek-van der Valk HJ, Wensing T, Breukink HJ. Weaning and the weanling diet influence the villous height and crypt depth in the small intestine of pigs and alter the concentrations of short-chain fatty acids in the large intestine and blood. J Nutr. 1998;128(6): 947–953.

42. Spreeuwenberg MA, Verdonk JM, Gaskins HR, Verstegen MW. Small intestine epithelial barrier function is compromised in pigs with low feed intake at weaning. J Nutr. 2001; 131(5): 1520–1527.

43. Montagne L, Pluske JR, Hampson DJ. A review of interactions between dietary fibre and the intestinal mucosa, and their consequences on digestive health in young non-ruminant animals. Anim Feed Sci Technol. 2003;108(1–4): 95–117.

44. Budiño FEL, Thomaz MC, Kronka RN, Nakaghi LSO, Tucci FM, Fraga AL, et al. Effect of probiotic and prebiotic inclusion in weaned piglet diets on structure and ultra-structure of small intestine. Brazilian Arch Biol Technol. 2005;48(6): 921–929.

45. Hedemann MS, Eskildsen M, Lærke HN, Pedersen C, Lindberg JE, Laurinen P, et al. Intestinal morphology and enzymatic activity in newly weaned pigs fed contrasting fiber concentrations and fiber properties. J Anim Sci. 2006;84(6): 1375–1386.

46. Awad WA, Ghareeb K, Abdel-Raheem S, Bohm J. Effects of dietary inclusion of probiotic and synbiotic on growth performance, organ weights, and intestinal histomorphology of broiler chickens. Poult Sci. 2009;88(1): 49–56.

47. Xu ZR, Hu CH, Xia MS, Zhan XA, Wang MQ. Effects of dietary fructooligosaccharide on digestive enzyme activities, and intestinal microflora and morphology of male broilers. Poult Sci. 2003;82: 1030–1036.

48. Buddle JR, Bolton JR. The pathophysiology of diarrhoea in pigs. Pig News Inf. 1992;13: 41–45.

49. Burrin DG, Stoll B, Van Goudoever JB, Reeds PJ. 1 9 Nutrient Requirements for Intestinal Growth and Metabolism in the Developing Pig. In: Digestive Physiology of Pigs: Proceedings of the 8th Symposium. CABI; 2001. p. 75.

50. Kisielinski K, Willis S, Prescher a, Klosterhalfen B, Schumpelick V. A simple new method to calculate small intestine absorptive surface in the rat. Clin Exp Med. 2002;2(3): 131–135.

51. Piel C, Montagne L, Sève B, Lallès J-P. Increasing digesta viscosity using carboxymethylcellulose in weaned piglets stimulates ileal goblet cell numbers and maturation. J Nutr. 2005;135(1): 86–91.

52. Wadolkowski EA, Laux DC, Cohen PS. Colonization of the Streptomycin-Treated Mouse Large-Intestine by A Human Fecal Escherichia-Coli Strain - Role of Growth in Mucus. Infect Immun. 1988;56(5): 1030–1035.

53. Edelman S, Leskelä S, Ron E, Apajalahti J, Korhonen TK. In vitro adhesion of an avian pathogenic Escherichia coli O78 strain to surfaces of the chicken intestinal tract and to ileal mucus. Vet Microbiol. 2003;91(1): 41–56.

54. Blomberg L, Krivan HC, Cohen PS, Conway PL. Piglet ileal mucus contains protein and glycolipid (galactosylceramide) receptors specific for Escherichia coli K88 fimbriae. Infect Immun. 1993;61(6): 2526–2531.

55. Blomberg L, Gustafsson L, Cohen PS, Conway PL, Blomberg A. Growth of Escherichia coli K88 in piglet ileal mucus: Protein expression as an indicator of type of metabolism. J Bacteriol. 1995;177(23): 6695–6703.

56. Pestova MI, Clift RE, Jason Vickers R, Franklin MA, Mathew AG. Effect of weaning and dietary galactose supplementation on digesta glycoproteins in pigs. J Sci Food Agric. 2000;80(13): 1918–1924.

57. Mathew A. Seeking Alternatives to Growth Promoting Antibiotics Alan. Manitoba Swine Semin. 2002;16: 115–128.

58. Erdem AL, Avelino F, Xicohtencatl-Cortes J, Girón JA. Host protein binding and adhesive properties of H6 and H7 flagella of attaching and effacing Escherichia coli. J Bacteriol. 2007;189(20): 7426–7435.

59. Bai X, Liu X, Su Y. Inhibitory effects of intestinal mucus on bacterial adherence to cultured intestinal epithelial cells after surface burns. Chin Med J. 2000;113(5): 449–450.

60. Korhonen J. Antibiotic Resistance of Lactic Acid Bacteria. PhD Thesis. 2010. 1–75 p.

61. Dowarah R, Verma AK, Agarwal N. The use of Lactobacillus as an alternative of antibiotic growth promoters in pigs: A review. Anim Nutr. 2017;3(1): 1–6.

62. Liao SF, Nyachoti M. Using probiotics to improve swine gut health and nutrient utilization. Anim Nutr. 2017;3(4): 331–343.

63. He P, Young LG, Forsberg C. Microbial transformation of deoxynivalenol (vomitoxin). Appl Environ Microbiol. 1992;58(12): 3857–3863.

64. Kollarczik B, Gareis M, Hanelt M. In Vitro Transformation of the Fusarium Mycotoxins Deoxynivafenof and Zearafenone by the Normal Gut Microflora of Pigs Birgit. Nat Toxins. 1994;2: 105–110.

65. Eriksen GS, Pettersson H, Johnsen K, Lindberg JE. Transformation of trichothecenes in ileal digesta and faeces from pigs. Arch Tierernahr. 2002;56(4): 263–274.

66. Niderkorn V, Boudra H, Morgavi DP. Binding of Fusarium mycotoxins by fermentative bacteria in vitro. J Appl Microbiol. 2006;101(4): 849–856.

67. Young JC, Zhou T, Yu H, Zhu H, Gong J. Degradation of trichothecenemycotoxins by chicken intestinal microbes. Food Chem Toxicol. 2007;45(1):136–43.

68. Sauer WC, Mosenthin R, Hartog L a Den. The effect of source of fiber on ilealand fecal amino acid digestibility and bacterial nitrogen excretion in growingpigs. J Anim Sci. 1991;69: 4070–4077.

69. Knudsen KEB, Jensen BB, Hansen I. Digestion of polysaccharides and othermajor components in the small and large intestine of pigs fed on diets consistingof oat fractions rich in β-D-glucan. Br J Nutr. 1993;70(2): 537–556.

70. Freire JPB, Guerreiro AJG, Cunha LF, Aumaitre A. Effect of dietary fibre sourceon total tract digestibility, caecum volatile fatty acids and digestive transit time inthe weaned piglet. Anim Feed Sci Technol. 2000;87(1–2): 71–83.

71. D’Aldebert E, Biyeyeme Bi Mve MJ, Mergey M, Wendum D, Firrincieli D,Coilly A, et al. Bile Salts Control the Antimicrobial Peptide Cathelicidin ThroughNuclear Receptors in the Human Biliary Epithelium. Gastroenterology. 2009;136(4): 1435–1443.

72. Nie Y, Hu J, Yan X. Cross-talk between bile acids and intestinal microbiota inhost metabolism and health. J Zhejiang Univ B. 2015;16(6): 436–446.

73. Burrin D, Stoll B, Moore D. Digestive physiology of the pig symposium:Intestinal bile acid sensing is linked to key endocrine and metabolic signalingpathways. J Anim Sci. 2013;91(5): 1991–2000.

74. Cremers CM, Knoefler D, Vitvitsky V, Banerjee R, Jakob U. Bile salts act aseffective protein-unfolding agents and instigators of disulfide stress in vivo. Proc Natl Acad Sci. 2014;111(16): E1610–E1619.

75. Kandell RL, Bernstein C. Bile salt/acid induction of DNA damage in bacterialand mammalian cells: implications for colon cancer. Nutr Cancer. 1991; 16(3–4):227–238.

76. Bernstein H, Payne CM, Bernstein C, Schneider J, Beard SE, Crowley CL. Activation of the promoters of genes associated with DNA damage, oxidative stress, ER stress and protein malfolding by the bile salt, deoxycholate. Toxicol Lett. 1999; 108(1): 37–46.

77. Bernstein C, Bernstein H, Payne CM, Beard SE, Schneider J. Bile salt activationof stress response promoters in Escherichia coli. Curr Microbiol. 1999;39(2): 68–72.

78. Chou JH, Greenberg JT, Demple B. Posttranscriptional repression of Escherichiacoli OmpF protein in response to redox stress: positive control of the micFantisense RNA by the soxRS locus. J Bacteriol. 1993;175(4): 1026–1031.

79. Oh JT, Cajal Y, Skowronska EM, Belkin S, Chen J, Van Dyk TK, et al. Cationic peptide antimicrobials induce selective transcription of micF and osmY in Escherichia coli. Biochim Biophys Acta. 2000;1463(1): 43–54.

80. Begley M, Gahan CGM, Hill C. The interaction between bacteria and bile. FEMS Microbiol Rev. 2005;29(4): 625–651.

81. Bederska-Lojewska D, Pieszka M. Modulating gastrointestinal microflora of pigs through nutrition using feed additives. Ann Anim Sci. 2011;11(3): 333–355.

82. Steer T.; Carpenter H.; Tuohy K.; Gibson G.R. Perspectives on the role of the human gut microbiota and its modulation by pro- and prebiotics. Nutr Res Rev. 2000;13(2000): 229–254.

83. Samanta AK, Jayapal N, Senani S, Kolte AP, Sridhar M. Prebiotic inulin: Useful dietary adjuncts to manipulate the livestock gut microflora. Brazilian J Microbiol. 2013;44(1): 1–14.

84. Blottière HM, Buecher B, Galmiche J-P, Cherbut C. Molecular analysis of the effect of short-chain fatty acids on intestinal cell proliferation. Proc Natl Acad Sci. 2003;62: 101–106.

85. Levrat M-A, Rémésy C, Demigné C. High propionic acid fermentations and mineral accumulation in the cecum of rats adapted to different levels of inulin. J Nutr. 1991;121(11): 1730–1737.

86. Campbell JM, Fahey GC, Wolf BW. Nutrient Metabolism Selected Indigestible Oligosaccharides Affect Large Bowel Mass, Cecal and Fecal Short-Chain Fatty Acids, pH and Microflora in Rats 1,2. J Nutr. 1997;127: 130–136.

87. Kleessen B, Hartmann L, Blaut M. Oligofructose and long-chain inulin: influenceon the gut microbial ecology of rats associated with a human faecal flora. Br J Nutr. 2001;86(2): 291–300.

88. Poulsen M, Mølck A, Jacobsen BL. Different effects of short- and long-chained fructans on large intestinal physiology and carcinogen-induced aberrant cryptfoci in rats. Nutr Cancer. 2002;42: 194–205.

89. Gibson GR, Probert HM, Loo J Van, Rastall RA, Roberfroid MB. Dietary modulation of the human colonic microbiota: updating the concept of prebiotics. Nutr Res Rev. 2004;17(2): 259–275.

90. Loh G, Eberhard M, Brunner RM, Hennig U, Kuhla S, Kleessen B, et al. Inulinalters the intestinal microbiota and short-chain fatty acid concentrations in growing pigs regardless of their basal diet. J Nutr. 2006;136(5): 1198–1202.

91. Barcenilla A, Pryde SE, Martin JC, Duncan SH, Stewart CS, Henderson C, et al. Phylogenetic relationships of butyrate-producing bacteria from the human gut. Appl Environ Microbiol. 2000;66(4): 1654–1661.

92. Duncan SH, Holtrop G, Lobley GE, Calder AG, Stewart CS, Flint HJ.Contribution of acetate to butyrate formation by human faecal bacteria. Br J Nutr. 2004;91(6):915–923.

93. Mølbak L, Thomsen LE, Jensen TK, Bach Knudsen KE, Boye M. Increased amount of Bifidobacterium thermacidophilum and Megasphaera elsdenii in the colonic microbiota of pigs fed a swine dysentery preventive diet containing chicory roots and sweet lupine. J Appl Microbiol. 2007;103(5): 1853–1867.

94. Patterson JK, Yasuda K, Welch RM, Miller DD, Lei XG. Supplemental Dietary Inulin of Variable Chain Lengths Alters Intestinal Bacterial Populations in Young Pigs. J Nutr. 2010;140(12): 2158–2161.

95. Liu H, Ivarsson E, Dicksved J, Lundh T, Lindberg JE. Inclusion of Chicory (Cichorium intybus L.) in pigs’ diets affects the intestinal microenvironment and the gut microbiota. Appl Environ Microbiol. 2012;78(12): 4102–4109.

96. Pérez D, Soraci A, Tapia M. Pharmacokinetics and bioavailability of calcium fosfomycin in post weaning piglets after oral administration. Int J Agro Vet Med Sci. 2012;6(6): 424–435.

97. Bernier SP, Surette MG. Concentration-dependent activity of antibiotics in natural environments. Front Microbiol. 2013;4: 1–14.

98. Munita JM, Arias CA, Unit AR, Santiago A De. HHS Public Access. Mech Antibiot Resist. 2016;4(2): 1–37.

99. Kahan FM, Kahan JS, Cassidy PJ, Kropp H. the Mechanism of Action of Fosfomycin (Phosphonomycin). Ann N Y Acad Sci. 1974;235(1): 364–386.

100. Gobernado M. Revisión Fosfomicina. Rev Española Quimioter. 2003;16(1): 15–40.

101. Castañeda-García A, Rodríguez-Rojas A, Guelfo JR, Blázquez J. The glycerol-3-phosphate permease GlpT is the only fosfomycin transporter in Pseudomonas aeruginosa. J Bacteriol. 2009;191(22): 6968–6974.

102. Popovic M, Steinort D, Pillai S, Joukhadar C. Fosfomycin: an old, new friend? Eur J Clin Microbiol Infect Dis. 2010;29(2): 127–142.

103. Pérez DS, Tapia MO, Soraci AL. Fosfomycin: Uses and potentialities in veterinary medicine. open Vet J. 2014;4(1): 26–43.

104. Nguyen LT, Haney EF, Vogel HJ. The expanding scope of antimicrobial peptide structures and their modes of action. Trends Biotechnol. 2011;29(9): 464–472.

105. Grenier B, Applegate TJ. Modulation of intestinal functions following mycotoxin ingestion: Meta-analysis of published experiments in animals. Toxins. 2013;5(2): 396–430.

106. Obremski K, Zielonka L, Gajecka M, Jakimiuk E, Bakula T, Baranowski M, et al. Histological estimation of the small intestine wall after administration of feed containing deoxynivalenol, T-2 toxin and zearalenone in the pig. Pol J Vet Sci. 2008;11(4): 339–345.

107. Bracarense A-PFL, Lucioli J, Grenier B, Drociunas Pacheco G, Moll W-D, Schatzmayr G, et al. Chronic ingestion of deoxynivalenol and fumonisin, alone or in interaction, induces morphological and immunological changes in the intestine of piglets. Br J Nutr. 2012;107(12): 1776–1786.

108. Gerez JR, Pinton P, Callu P, Grosjean F, Oswald IP, Bracarense AP aula FL. Deoxynivalenol alone or in combination with nivalenol and zearalenone induce systemic histological changes in pigs. Exp Toxicol Pathol. 2015;67(2): 89–98.

109. Laparra JM, Sanz Y. Comparison of in vitro models to study bacterial adhesion to the intestinal epithelium. Lett Appl Microbiol. 2009;49(6): 695–701.

110. Pinton P, Graziani F, Pujol A, Nicoletti C, Paris O, Ernouf P, et al. Deoxynivalenol inhibits the expression by goblet cells of intestinal mucins through a PKR and MAP kinase dependent repression of the resistin-like molecule β. Mol Nutr Food Res. 2015;59(6): 1076–1087.

111. Robert H, Payros D, Pinton P, Théodorou V, Mercier-Bonin M, Oswald IP. Impact of mycotoxins on the intestine: are mucus and microbiota new targets? J Toxicol Environ Heal - Part B Crit Rev. 2017;20(5): 249–275.

112. Pérez Gaudio DS, Martínez G, Fernández Paggi MB, Riccio MB, Decundo JM, Diéguez S, et al. Protective Effect of Fosfomycin on Deoxynivalenol- Treated Cell Cultures. Eur J Biomed Pharm Sci. 2016;3(7): 99–106.

113. Niewold TA. The nonantibiotic anti-inflammatory effect of antimicrobial growth promoters, the real mode of action? A hypothesis. Poult Sci. 2007;86:605–609.

114. Costa E, Uwiera RR, Kastelic JP, Selinger LB, Inglis GD. Non-therapeutic administration of a model antimicrobial growth promoter modulates intestinal immune responses. Gut Pathog. 2011;3(1): 1–15.

115. Niewold TA. Mechanisms of antibiotics: how do they really work. Banff Pork Semin Proceedings, Banff, Alberta, Canada, 15-17 January 2013. 2013;24(2013): 103–106.

116. Brown K, Zaytsoff SJM, Uwiera RRE, Inglis GD. Antimicrobial growth promoters modulate host responses in mice with a defined intestinal microbiota. Sci Rep. 2016;6: 1–13.

117. Brown K, Uwiera RRE, Kalmokoff ML, Brooks SPJ, Inglis GD. Antimicrobial growth promoter use in livestock: a requirement to understand their modes of action to develop effective alternatives. Int J Antimicrob Agents. 2017; 49(1): 12–24.

118. Collington GK, Parker DS, Armstrong DG. The influence of inclusion of either an antibiotic or a probiotic in the diet on the development of digestive enzyme activity in the pig. Br J Nutr. 1990;64(1): 59–70.

119. Touchette KJ, Carroll JA, Allee GL, Matteri RL, Dyer CJ, Beausang LA, et al. Effect of spray-dried plasma and lipopolysaccharide exposure on weaned pigs: I. Effects on the immune axis of weaned pigs. K J Touchette, J A Carroll, G L Allee, R L Matteri, C J Dyer, L A Beausang and M E Zannelli The online version of this artic. J Anim Sci. 2002;80: 494–501.

120. Liu P, Piao XS, Kim SW, Wang L, Shen YB, Lee HS, et al. Effects of chito-oligosaccharide supplementation on the growth performance, nutrient digestibility, intestinal morphology, and fecal shedding of and in weaning pigs. J Anim Sci. 2008;86(10): 2609–2618.

121. le Roux CW, Borg C, Wallis K, Vincent RP, Bueter M, Goodlad R, et al. Gut Hypertrophy After Gastric Bypass Is Associated With Increased Glucagon-Like Peptide 2 and Intestinal Crypt Cell Proliferation. Ann Surg. 2010;252(1): 50–56.

122. Jain AK, Stoll B, Burrin DG, Holst JJ, Moore DD. Enteral bile acid treatment improves parenteral nutrition-related liver disease and intestinal mucosal atrophy in neonatal pigs. AJP Gastrointest Liver Physiol. 2012;302(2): G218–G224.

123. de Diego-Cabero N, Mereu A, Menoyo D, Holst JJ, Ipharraguerre IR. Bile acid mediated effects on gut integrity and performance of early-weaned piglets. BMC Vet Res. 2015;11(1): 1–8.

124. Stojancevic M, Stankov K, Mikov M. The impact of farnesoid X receptor activation on intestinal permeability in inflammatory bowel disease. Can J Gastroenterol. 2012;26(9): 631–637.

125. Morikawa K, Oseko F, Morikawa S, Sawada M. Immunosuppressive activity of fosfomycin on human T-lymphocyte function in vitro. Antimicrob Agents Chemother. 1993;37(12): 2684–2687.

126. Perez Fernandez P, Herrera I, Martinez P, Gomez-Lus ML, Prieto J. Enhancement of the susceptibility of Staphylococcus aureus to phagocytosis after treatment with fosfomycin compared with other antimicrobial agents. Chemotherapy. 1995;41(1): 45–49.

127. Krause R, Patruta S, Daxböck F, Fladerer P, Wenisch C. The effect of fosfomycin on neutrophil function. J Antimicrob Chemother. 2001;47(2): 141–146.

128. Morikawa K, Zhang J, Nonaka M, Morikawa S. Modulatory effect of macrolide antibiotics on the Th1- and Th2-type cytokine production. Int J Antimicrob Agents. 2002;19(1): 53–59.

129. Tullio V, Cuffini AM, Banche G, Mandras N, Allizond V, Roana J, et al. Role of fosfomycin tromethamine in modulating non-specific defence mechanisms in chronic uremic patients towards ESBL-producing Escherichia coli. Int JImmunopathol Pharmacol. 2008;21(1): 153–160.

130. Buret AG. Immuno-modulation and anti-inflammatory benefits of antibiotics:The example of tilmicosin. Can J Vet Res. 2010;74(1): 1–10.

131. Shen YB, Piao XS, Kim SW, Wang L, Liu P, Yoon I, et al. Effects of yeast culture supplementation on growth performance, intestinal health, and immuneresponse of nursery pigs. J Anim Sci. 2009;87(8): 2614–2624.

132. Yokota SI, Okabayashi T, Yoto Y, Hori T, Tsutsumi H, Fujii N. Fosfomycin suppresses RS-virus-induced Streptococcus pneumoniae and Haemophilus influenzae adhesion to respiratory epithelial cells via the platelet-activating factor receptor. FEMS Microbiol Lett. 2010;310(1): 84–90.

133. Michalopoulos AS, Livaditis IG, Gougoutas V. The revival of fosfomycin. Int JInfect Dis. 2011;15(11): e732–e739.

134. Zhang Y, Limaye PB, Renaud HJ, Klaassen CD. Effect of various antibiotics on modulation of intestinal microbiota and bile acid profile in mice. Toxicol Appl Pharmacol. 2014;277(2): 138–145.

135. Falagas ME, Vouloumanou EK, Samonis G, Vardakas KZ. Fosfomycin. ClinMicrobiol Rev. 2016;29(2): 321–347.

